# Representation Learning of Human Diseases for Indication Expansion and Investment Decisions

**DOI:** 10.1101/2024.11.19.624381

**Authors:** Cyrus Babak Ravandi, William R. Mowrey, Ayan Chatterjee, Fariba Khanshan, Parham Haddadi, Michaël Ughetto, Ian Barrett, Wei Ding, Guillermo Del Angel, Tom Diethe, Juan Carlos Mobarec, Simon Lambden, Seng Cheng, Tina Eliassi-Rad, Gabriel Risa, Piero Ricchiuto

## Abstract

A fundamental challenge in translational medicine is the computational modeling of complex human diseases to accelerate therapeutic development. Representation learning provides a powerful framework to address this, yet creating models that capture deep biological mechanisms remains a critical need. To this end, we propose a novel strategy that partitions the disease landscape into rare and non-rare categories, enabling systematic knowledge repurposing both within and between these groups. Here, we introduce Dis2Vec (Disease to Vector), a representation learning framework designed to operationalize this concept. Dis2Vec generates biologically grounded disease embeddings by learning from human genetic and phenotypic data, forming the foundation for Disease-Disease Association Learning (DDAL) and unsupervised disease clustering. We evaluate Dis2Vec representations in two downstream applications. First, we assess DDAL performance on a transfer learning benchmark designed to predict therapeutic transferability, using real-world drug repurposing investment decisions made in clinical trials. Second, unsupervised clustering analyses reveal shared biological mechanisms across diseases. By modeling the disease landscape in this way, Dis2Vec enhances translational research efficiency across both rare and non-rare diseases, advancing the development of foundational models for therapeutic science. Furthermore, Dis2Vec establishes a biologically grounded disease-representation and benchmarking layer that paves the way for trustworthy agentic biomedical AI systems in rare-disease indication expansion.

## Introduction

A fundamental challenge in translational medicine is the computational modeling of complex human diseases to accelerate therapeutic development [1, 2]. While representation learning offers a powerful framework for this task, a critical need remains for models that recover biologically meaningful mechanisms rather than relying on superficial statistical associations [3–5]. Current approaches primarily rely on Natural Language Processing (NLP) or Knowledge Graphs (KGs) [6, 7]. However, KGs face the challenge of aggregating noise and inductive biases from diverse data sources [8–10]. Recent advancements have introduced Generative AI agents, such as Biomni [11], that can actively reason through biological mechanisms. While these systems can generate useful hypotheses and orchestrate biomedical tools, their trustworthiness depends on access to biologically grounded representations and benchmarks that constrain mechanistic inference [12–16]. This motivates the need for disease representations that are not only predictive, but also grounded in shared mechanisms across the broader landscape of human disease.

Addressing this gap requires disease representations that capture shared biological mechanisms both within and across rare and non-rare disease contexts. Knowledge repurposing across these contexts offers a promising path toward a foundational model of disease pathogenesis, particularly because rare diseases span many therapeutic areas and can illuminate mechanisms that cut across traditional disease categories. Historically, rare disease research has enhanced our understanding of non-rare diseases; conversely, genetic signals from non-rare diseases can help complete the mechanistic picture for rare ones, where population studies are often unfeasible [17–22]. More broadly, disease-agnostic representations can support knowledge transfer both across and within disease categories. This multidirectional exchange is particularly valuable because rare diseases span a wide range of therapeutic areas, offering opportunities to identify shared biological mechanisms across conditions that are often studied in isolation. Embracing these “disease associations” (Figure 1A-B) provides a unique opportunity to develop comprehensive representations of diseases and their relationships. By clustering diseases based on shared biological mechanisms, we can address the challenge of data siloing [23–28] and accelerate the development of new therapies [29].

**Figure 1:**
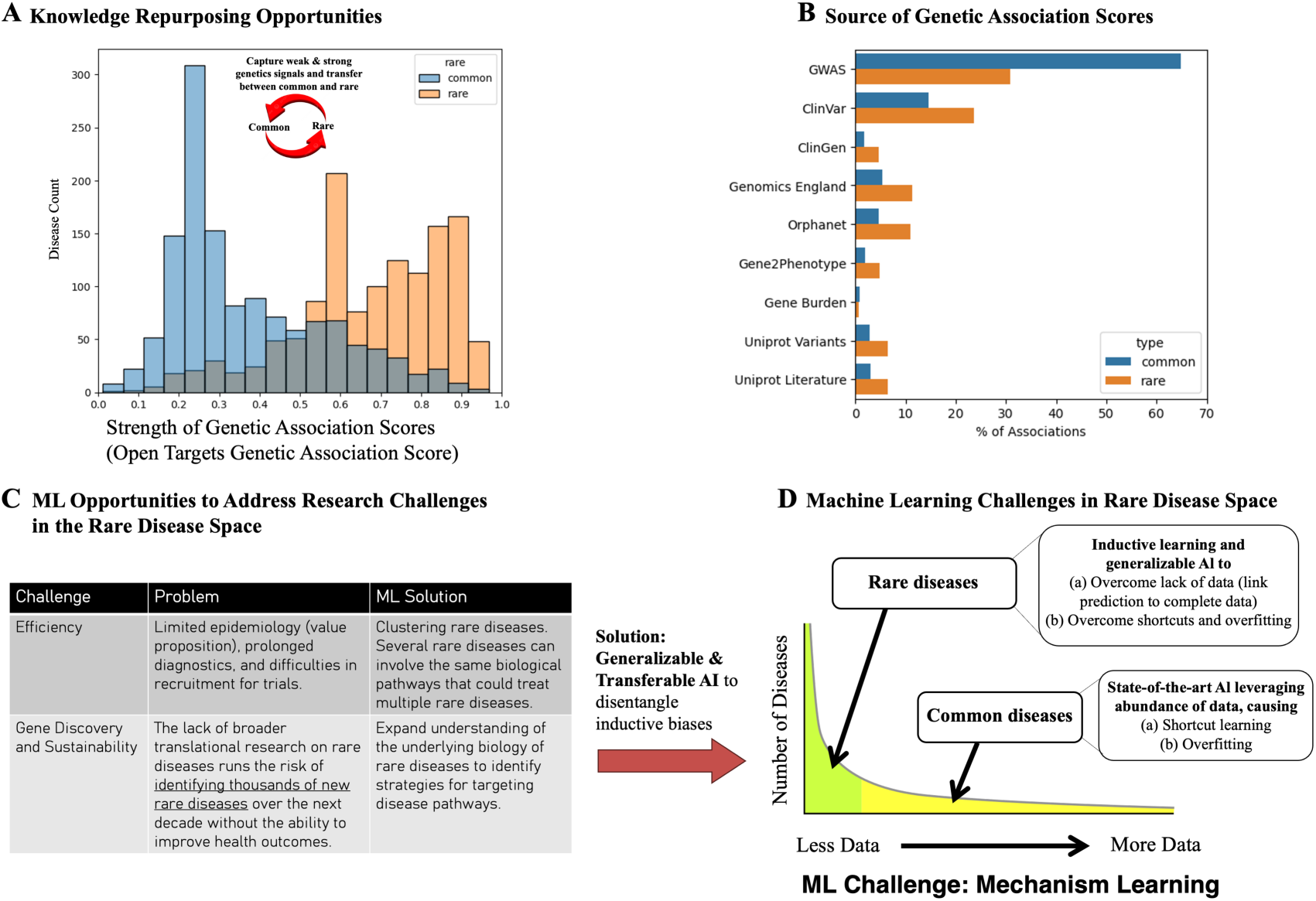
Repurposing Opportunities for ML to address Research Challenges in the Rare and non-rare Diseases. **(A-B)** Both rare and non-rare diseases are driven by Human biology. Rare diseases have a stronger genetic association score (obtained from OpenTargets [30] that combines various databases), driven by strong effect genetic variants that are causal for diseases. In contrast, non-rare diseases have a weaker genetic association score, driven by GWAS which has limited power to uncover causal links. Hence there is a knowledge repurposing opportunity to capture weak and strong genetics signals and transfer between non-rare and rare diseases. **(C-D)** Generalizable and transferable ML offers opportunities to address efficiency and discovery challenges in the data scarce regime.

From a machine learning perspective, this strategy is particularly valuable because rare diseases force models to operate in a data-scarce regime (Figure 1C-D). This scarcity necessitates more stringent and realistic evaluations, such as transfer learning [31, 32] and inductive tests [33–35], which compel a model to learn the underlying biological mechanisms rather than simply memorizing patterns in abundant data. This approach helps avoid common pitfalls in data-rich environments, like shortcut learning and class imbalance [33, 36, 37], which can artificially inflate performance and hinder the generalizability of state-of-the-art models.

In this paper, we introduce Dis2Vec, a disease representation learning framework that maps both rare and non-rare diseases into a shared continuous embedding space. We validate the quality of these embeddings through two challenging downstream tasks: Disease-Disease Association Learning (DDAL) and unsupervised disease clustering. For the DDAL task, we use the Dis2Vec embeddings to develop a two-shot learning architecture [38], UPNA-DDAL (Unsupervised Pre-training of Node Attributes for DDAL), and evaluate its performance on predicting drug repurposing decisions from clinical trials. For the clustering task, we demonstrate the ability of these embeddings to group diseases based on shared biological pathways. Ultimately, these results suggest that the high-quality representations from Dis2Vec can serve as a valuable resource for developing the generalizable and foundation models needed to advance therapeutic science.

## Results

### Disease to Vector

To generate meaningful representations of diseases, particularly for those with sparse data, a model must capture the complex higher-order relationships between underlying biological processes and observable symptoms. A significant challenge is that many diseases lack comprehensive annotations; for example, 53% of diseases have either no discovered genetic associations or no curated symptoms in Orphanet (see Methods and Figure S2A). This data incompleteness requires a method that can effectively combine and repurpose knowledge from multiple sources to learn the mechanisms of action.

To address this, we created the Dis2Vec representation learning pipeline, which builds a novel disease-disease network and then learns vector embeddings from its structure (Figure 2). The pipeline’s primary innovation is the creation of the MechDist (Ontology Distance) network (see Methods and Figure S4). Unlike standard knowledge graphs that often rely on simple linear associations, MechDist is built by computing the mechanistic similarity between every pair of diseases. This is achieved by minimizing the Lowest Common Ancestor (LCA) distances for each disease pair within the rich, hierarchical structures of the Gene Ontology (GO) and Human Phenotype Ontology (HPO). Once the MechDist network is constructed, we use Node2Vec [39] to generate the final Dis2Vec embeddings, as its algorithm is adept at learning higher-order dependencies by exploring both local and distant neighborhoods in the graph.

**Figure 2:**
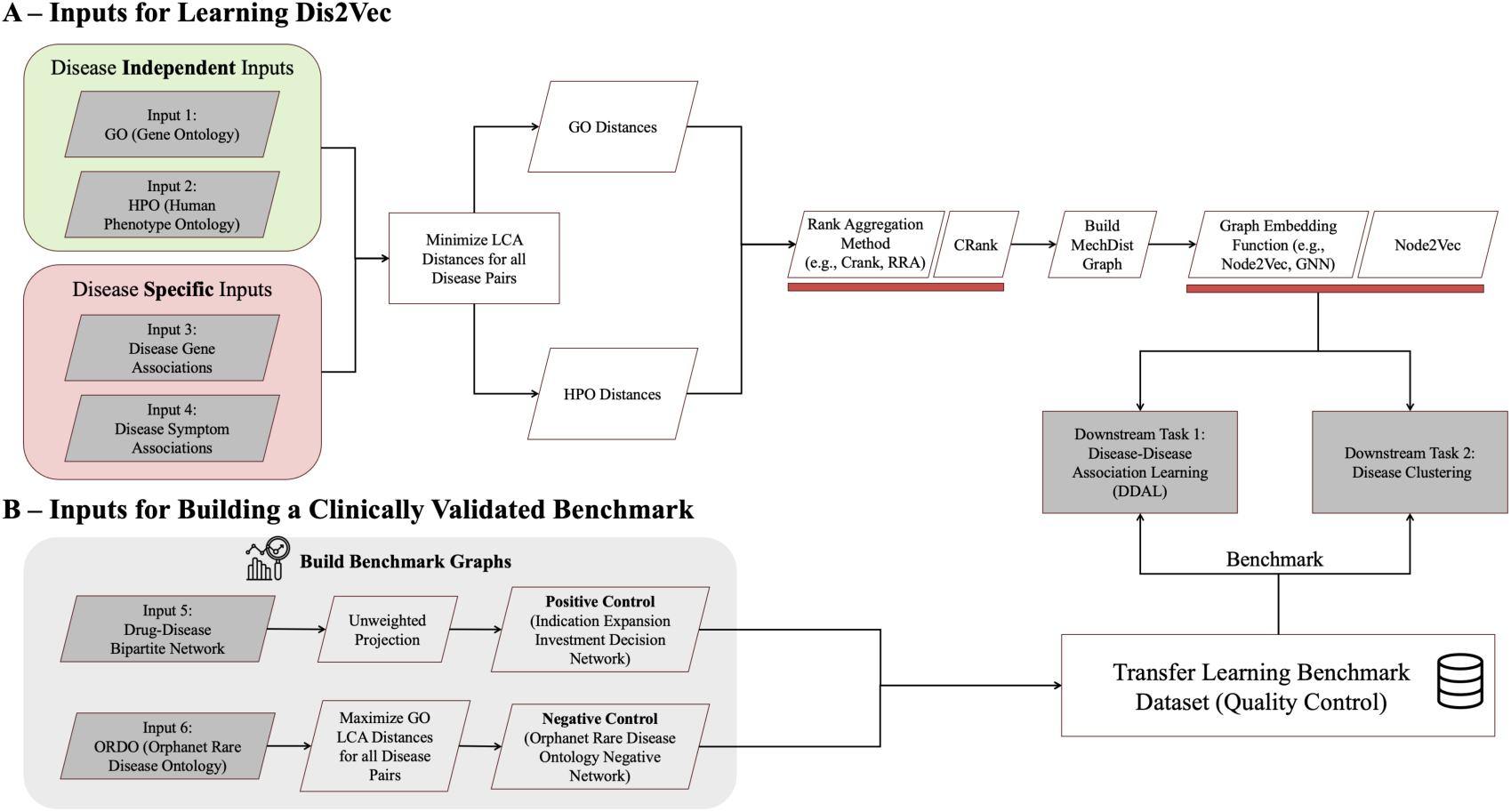
Dis2Vec Pipeline. **(A)** Dis2Vec is a multi-modal representation learning pipeline, enabling to build a disease-disease association network, namely MechDist, by combining biological knowledge from various sources. We incorporate higher-order knowledge from ontologies by minimizing the LCA distances for all diseases pairs (see Methods). Next, we used CRank to aggregate and prioritize LCA distances due to its ability to handle partial data, favor top-tier consensus, and maintain network-topology awareness. Lastly, any graph embedding function capable of aggregating both local (breadth first) and far neighborhoods (depth first) can be used on MechDist network to obtain Dis2Vec (we used Node2Vec). **(B)** We construct positive and negative controls utilizing independent data modalities not used in learning Dis2Vec embeddings [40]. For positive controls, we utilize drug repurposing investment decisions from clinical trials, obtaining disease-disease relationships where two diseases entered trials with the same drug. For negative controls, we used ORDO by maximizing LCA distances to identify disease pairs that are significantly different in terms of underlying biology.

Finally, a critical component of our framework is a robust benchmarking strategy designed to validate the biological relevance of the Dis2Vec embeddings. We evaluate the learned representations using data modalities that are entirely independent of the inputs used to create them (Figure 2B). This ensures that the embeddings capture true biological signals rather than simply memorizing features from the training data.

### Benchmarking Disease-Disease Association Learning

To properly evaluate a model’s ability to predict Disease-Disease Associations (DDA), we utilize our cross-evidence and robust benchmark networks [40], designed to test generalizability for the DDAL task. A core principle of our cross-evidence benchmarks is the separation of training and evaluation data. While information on drug targets is suitable for testing a foundational model of disease pathogenesis, it is not appropriate for training one because it is highly biased; only about 10% of druggable human proteins are ever targeted in clinical trials, representing a very small fraction of human biology [41]. This cross-evidence benchmark, consists of distinct positive and negative controls derived from sources independent of Dis2Vec training data. The positive controls are based on real-world investment decisions from the pharmaceutical industry, namely Indication Expansion Investment Decision Network (IxIDN) from a drug-disease bipartite network derived from clinical trials [40]. In this network, an edge connects two diseases if they have both been targets of a clinical trial for the same drug, signifying a strong, financially-backed hypothesis of a shared biological mechanism. Similarly, negative controls are needed to ensure the model can correctly identify and reject unrelated diseases. We used the Orphanet Rare Disease Ontology Negative-network (ORDON) introduced in [40], which is obtained by maximizing LCA distances in ORDO hierarchy [42].

Remarkably, even before using IxIDN and ORDON in a downstream predictive task, the Dis2Vec embeddings inherently capture the signals in our benchmark. A simple calculation of the Euclidean distance between disease vectors clearly separates the positive controls from the IxIDN and the negative controls from the ORDON (Figure 3C-D).

**Figure 3:**
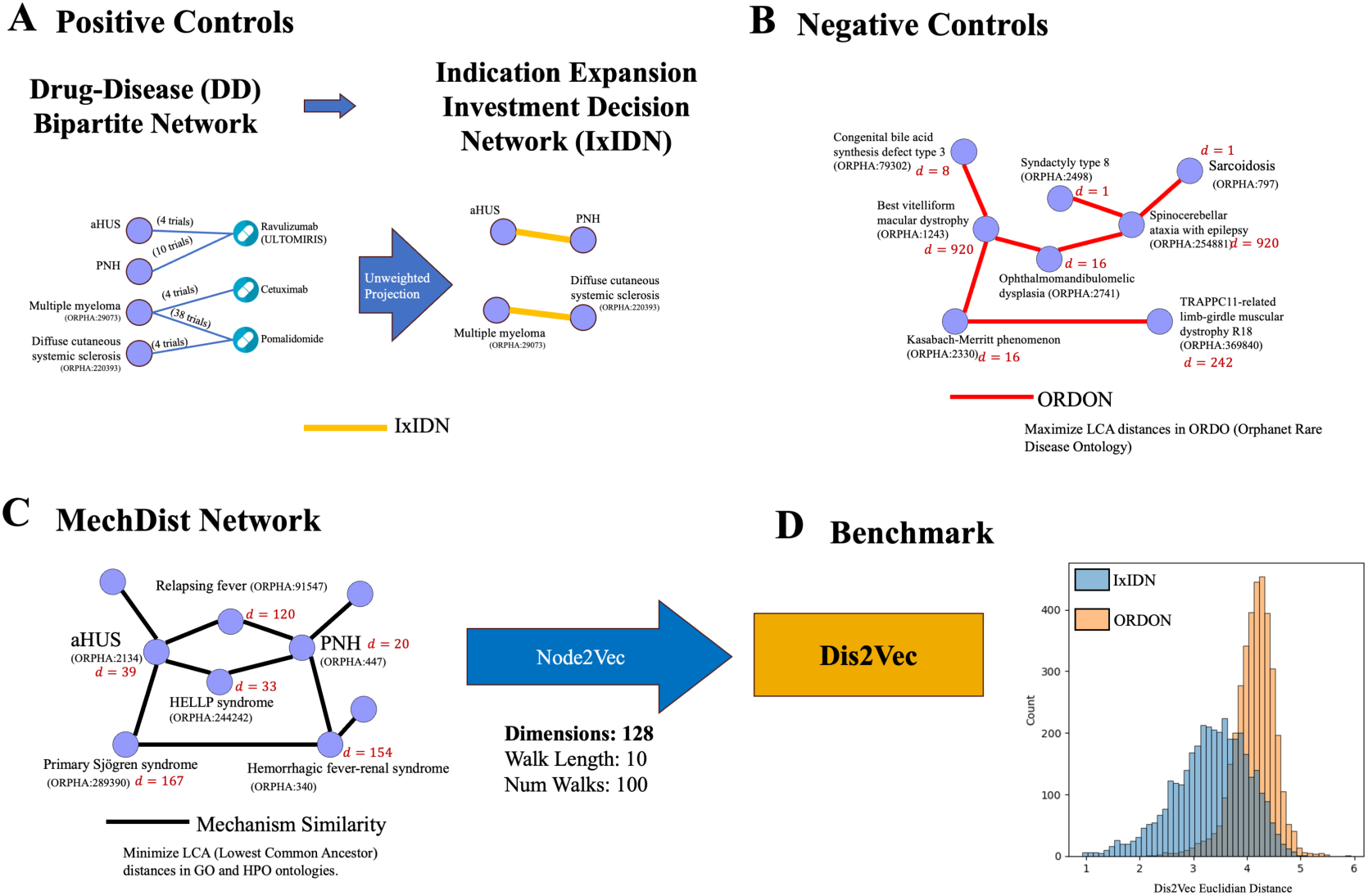
Transfer Learning Benchmark and Disease Representation Learning (Disease to Vector). **(A-B)** We establish transfer learning benchmark by utilizing IxIDN [40] derived from drug repurposing investment decisions made in clinical trials as positive samples. For negative samples, we established an orthogonal transfer learning benchmark by using ORDON [40] based on maximizing LCA distances in Orphanet Rare Disease Ontology. **(C)** We created MechDist network by combining non-linear dependencies in ontologies (GO and HPO) with disease genetics and symptoms associations (Methods). MechDist is an unweighted network where an edge represents two disease share biology according to the knowledge enriched in GO and HPO. **(D)** We observe that indeed MechDist’s structure captures higher order biological relationships between diseases by utilizing Node2Vec to learn disease representation. Remarkably, the Euclidean distance of Dis2Vec embeddings distinguish the positive and negative transfer learning benchmarks, demonstrating excellent classification capabilities obtained from rich disease representations that can be utilized for downstream ML tasks such as disease association learning and clustering.

### Validating the Dis2Vec Representations via Downstream Tasks

To validate the utility of our Dis2Vec embeddings, we designed two downstream tasks that reflect key challenges in translational medicine. These tasks test the coherence of the learned representations at two different scales: the pairwise level of individual disease-disease relationships and the higher-order level of the global disease landscape. A truly useful disease representation model must excel at both tasks:

- **[Downstream Task 1 - Link Prediction] Disease-Disease Association Prediction:** This link prediction task tests whether the embeddings can effectively model the relationship between pairs of diseases. We demonstrate that the pre-trained Dis2Vec representations can be used by a link prediction decoder to forecast drug-target investment decisions in a rigorous transfer learning setting.
- **[Downstream Task 2 - Community Detection] Unsupervised Disease Clustering:** This node labeling task addresses a major source of inefficiency in medical research: the artificial separation of diseases by clinical specialty. We show that instead of treating diseases as isolated entities, the Dis2Vec embeddings can be used to automatically cluster them based on shared mechanisms of action.

We will now assess the performance and biological interpretability of Dis2Vec on these two tasks.

### Dis2Vec Demonstrates High Performance on Disease-Disease Association Prediction

To evaluate the first downstream task, we assessed the capacity of Dis2Vec embeddings to accurately predict real-world drug repurposing decisions. We implemented a rigorous transfer learning framework to ensure the model generalizes from underlying biological principles rather than simply exploiting topological shortcuts or memorizing a drug–disease graph structures [35]. First, we trained a link prediction decoder, named UPNA-DDAL, using the Dis2Vec embeddings (Figure 4). The model was trained exclusively on the MechDist network, which captures fundamental biological and phenotypic similarities between diseases. Critically, all positive and negative edges from our benchmark datasets, IxIDN and ORDON, were removed from the training data. The trained model was then tested on its ability to reconstruct IxIDN and ORDON.

**Figure 4:**
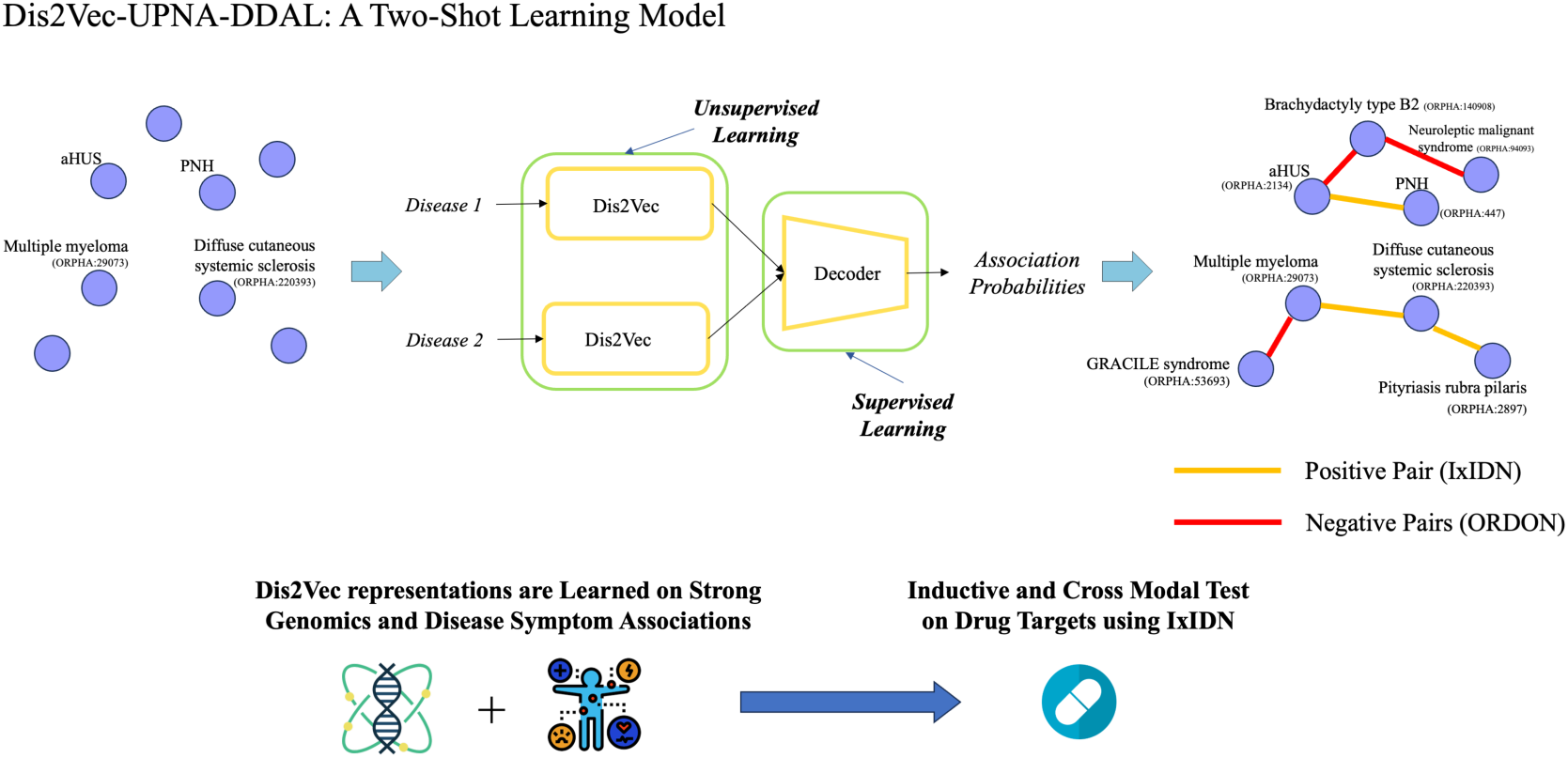
Generalizable Model for Disease-Disease Association Learning. **(A)** The first downstream task for Dis2Vec is DDAL, for which we train UPNA-DDAL, a two-shot learning model [38], to decode the biological signals captured by Dis2Vec. Dis2Vec learns disease representations from human genetic evidence and observed symptoms. We formulate DDAL as a transfer learning task using IxIDN and ORDON as positive and negative samples, respectively. IxIDN is derived from clinical trials, which often do not directly target causal genetic links, whereas ORDON is derived from the Orphanet Rare Disease Ontology hierarchy [40].

The model showed strong predictive power in this transfer learning scenario, achieving an Area Under the Receiver Operating Characteristic Curve (AUROC) of 0.89. This high performance indicates that the biological signals encoded in the Dis2Vec embeddings effectively capture the complex reasoning behind pharmaceutical investment decisions. As shown in Figure 5B, the model’s predicted association probabilities clearly separate the positive investment decisions from the ontologically distant negative controls.

**Figure 5:**
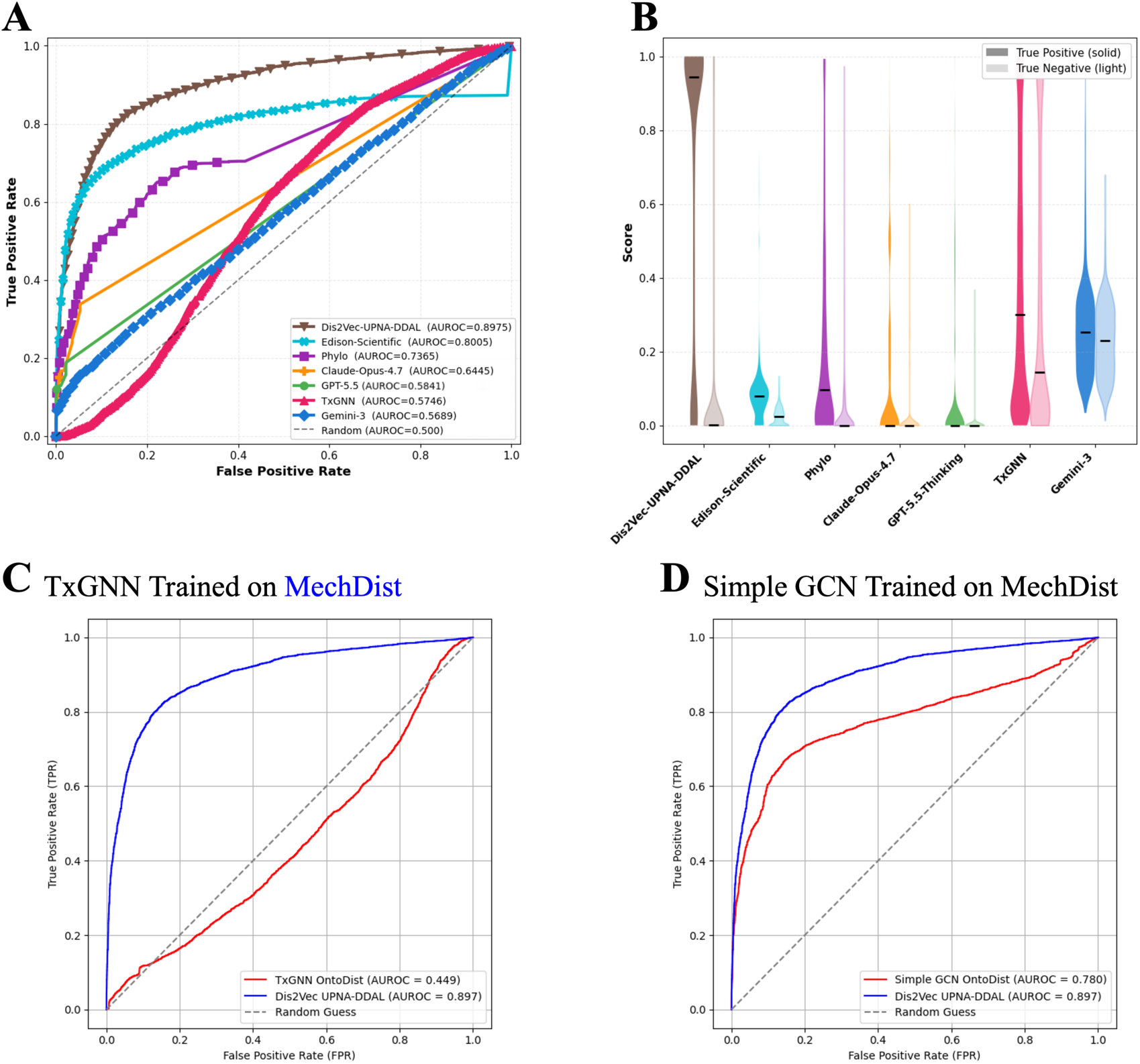
Cross-evidence Generalization Benchmark on IxIDN and ORDON. **(A) ROC curves.** Performance of GraphML models and agentic/ LLM systems on the IxIDN/ORDON benchmarks [40] under cross-evidence generalization protocols. GraphML models consist of TxGNN trained on heterogeneous PrimeKG with drugs removed to control shortcuts and Dis2Vec-UPNA-DDAL on homogeneous MechDist network. All IxIDN and ORDON edges were removed from PrimeKG and MechDist during training (Protocol 1, <u>see</u> Methods). LLMs (GPT-5.5-Thinking, Claude-Opus-4.7, and Gemini-3) and Agentic AI systems (Phylo and Edison Scientific) were evaluated via structured prompting on blinded disease pairs (Protocol 2; Methods). **(B) Score distributions.** Violin and box plots of predicted DDA scores for true-positive IxIDN pairs versus true-negative ORDON pairs. **(C)** To evaluate PrimeKG’s heterogeneity as the sole cause of TxGNN performance degradation under drug-node removal and held-out benchmark conditions, we modified TxGNN relational meta-path architecture and trained TxGNN on MechDist. However, **(D)** shows that a simple Graph Convolutional Neural Networks (GCN) outperforms the modified TxGNN on DDAL task over MechDist. This indicates that a homogeneous MechDist graph and two-shot learning architecture is a better fit for DDAL task vs heterogeneous KG with zero-shot learning.

### Case Study: Decoding Shared Mechanisms

A compelling example of the model’s capability is its confident prediction of the link between atypical hemolytic uremic syndrome (aHUS) and paroxysmal nocturnal hemoglobinuria (PNH), two disorders treated with the FDA-approved drug Ravulizumab [43]. This prediction, presented in Figure 4 (score: 0.99), is particularly insightful because the two diseases are not directly connected in the MechDist training network and do not share causal genes.

### Case Study: Bridging Rare and Non-Rare Diseases

The model also excels at identifying repurposing opportunities from non-rare to rare diseases. For instance, UPNA-DDAL predicts strong associations between the non-rare autoimmune disorder Rheumatoid Arthritis (RA, EFO_0000685) and a group of rare Idiopathic Inflammatory Myopathies (IIMs) [44], including Antisynthetase Syndrome (ORPHA:81, score: 0.98). Antisynthetase Syndrome disorder presents joint manifestations that can, in some cases, be considered a rheumatoid arthritis overlap syndrome [45], and both disorders share the genetic risk associated with specific haplotypes in the HLA region [46]. Most other IIMs and overlap disorders also showed strong association scores for the disorders, including Polymyositis (ORPHA:732, score: 0.94), Dermatomyositis (ORPHA:645613, 0.94), Mixed Connective Tissue Disease (ORPHA:809, 0.98) and Overlap Myositis (ORPHA:206572, 0.97).

### Evaluating Dis2Vec as a Grounding Layer for LLM, Agentic AI, and GraphML Systems

To rigorously assess the generalization power of Dis2Vec relative to contemporary approaches, we evaluated four categories of systems under strict **cross-evidence generalization** protocols designed to prevent information leakage from training to test data (see Table 1). These protocols reflect the native capabilities and constraints of each model class while ensuring a fair, decision-oriented test using the held-out IxIDN and ORDON benchmarks.

- **Protocol 1 (GraphML models)**: Dis2Vec-UPNA-DDAL and a simple GCN were trained on the homogeneous MechDist network. TxGNN [47] was trained on the heterogeneous PrimeKG. In all cases, IxIDN and ORDON edges were completely removed from training, and drug nodes were excluded from TxGNN to eliminate shortcut learning via known drug–disease associations.
- **Protocol 2a (LLMs, parametric only)**: GPT-5.5 [48], Claude Opus 4.7 [49], and Gemini 3 [50] scored. In GraphML, edge removal provides a direct mechanism for preventing leakage and constructing an inductive evaluation setting. For LLMs, however, we cannot guarantee that the data sources underlying IxIDN and ORDON, namely clinical trials and ORDO, were excluded from model training. To reduce potential source leakage, we explicitly instructed the models not to use these sources, randomized disease pairs, and removed ontology IDs prior to evaluation. We did not restrict the models from using web search.
- **Protocol 2b (Agentic AI)**: Edison Scientific [51, 52] and Phylo [53] used the same prompt as Protocol 2a. The main difference from Protocol 2a is that these agentic AI tools are specialized for bioinformatics tasks and are equipped with MCPs that enable them to run bioinformatics tools.

**Table 1:** Summary of cross-evidence generalization protocols applied across system categories.

| Protocol | System type | Examples | Evaluation approach |
| --- | --- | --- | --- |
| 1 | GraphML on Homogeneous- / Heterogeneous -graphs (unipartite graph vs KG) | Dis2Vec/<br>UPNA-DDAL,<br>TxGNN | Retrain on MechDist with IxIDN + ORDON + drug nodes removed; test on held-out IxIDN/ORDON |
| 2a | LLM (parametric only) | GPT-5.5,<br>Claude-Opus-4.7,<br>Gemini-3 | Structured prompting; DDAL score from parametric knowledge only; no tool calls |
| 2b | Agentic AI (tool-augmented) | Phylo, Edison<br>Scientific | Same structured prompt; model may use tools or retrieval at inference time |

Figure 5A presents the results of this comprehensive evaluation, showing the ROC curves for all systems. Dis2Vec-UPNA-DDAL achieved the strongest performance under this cross-evidence, suggesting that biologically grounded disease representations capture information not reliably recovered by current general-purpose or agentic systems alone. Among the evaluated agentic systems, Edison Scientific performed best, achieving an AUROC of 0.8005, which exceeded the performance of the other LLM-based systems and the 0.55 to 0.68 AUROC range observed for most parametric and tool-augmented models. By comparison, TxGNN trained on PrimeKG reached only an AUROC of 0.575, indicating weaker generalization in this setting. This gap may reflect the dependence of pure knowledge graph models on fixed graph structure and observed edges, whereas full agentic systems can combine reasoning, retrieval, tool use, and broader contextual evidence during inference. This trend is consistent with the broader movement from standalone KG-based approaches toward agentic-augmented KG frameworks, where knowledge graphs are used as reasoning substrates rather than closed prediction systems [54]. Figure 5B further highlights these differences through violin and box plots of the predicted DDA scores. Dis2Vec-UPNA-DDAL produced a clear bimodal distribution with strong separation between IxIDN positives and ORDON negatives, while most other evaluated systems showed weaker separation and substantially overlapping score distributions under this benchmark. These results suggest that general-purpose reasoning, retrieval, and KG-based inference alone may not consistently recover rare-disease mechanistic transfer signals, although specialized agentic systems such as Edison Scientific can substantially narrow this gap.

To determine whether TxGNN’s substantially reduced performance was due to PrimeKG’s heterogeneity, we modified its relational meta-path architecture and retrained it on the homogeneous MechDist network (Figure 5C). Performance declined further to an AUROC of 0.449 (below random chance). In comparison, a simple Graph Convolutional Network (GCN) trained on the same MechDist graph achieved a substantially higher AUROC of 0.780 (Figure 5D). These results demonstrate that the combination of a homogeneous disease–disease graph and Dis2Vec’s two-shot learning architecture is better aligned with this DDAL benchmark, where the task requires disease-level mechanistic transfer rather than retrieval of known drug-disease paths.

Collectively, these findings show that most existing approaches exhibit limited discrimination under the demanding cross-evidence generalization setting required by IxIDN and ORDON. The results underscore the value of biologically grounded, mechanism-centered representations and decision-oriented benchmarks for advancing translational AI. Next, we assess the utility of IxIDN/ORDON in a new task, unsupervised disease clustering, using Dis2Vec framework.

### Unsupervised Clustering via Dis2Vec Reveals a Biologically Coherent Disease Landscape

For the second downstream task, we evaluated whether Dis2Vec embeddings could organize the complex landscape of human disease into a mechanistically coherent map. To achieve this, we performed unsupervised clustering using a Gaussian Mixture Model (GMM) over Dis2Vec 128-dimensional disease vectors, which accommodates non-spherical cluster geometries and enforces probabilistic soft assignments rather than rigid boundaries. The results, visualized in two dimensions with UMAP (Figure 6A), reveal a global structure with clear biological relevance. While some clusters are defined by anatomy-likely driven by shared phenotypes-many others group diseases by underlying biological processes, demonstrating that the model captures more than just superficial symptom similarity. A prime example is Cluster 0, which contains a diverse set of immune system dysregulations, including cell deficiencies and functional abnormalities.

**Figure 6:**
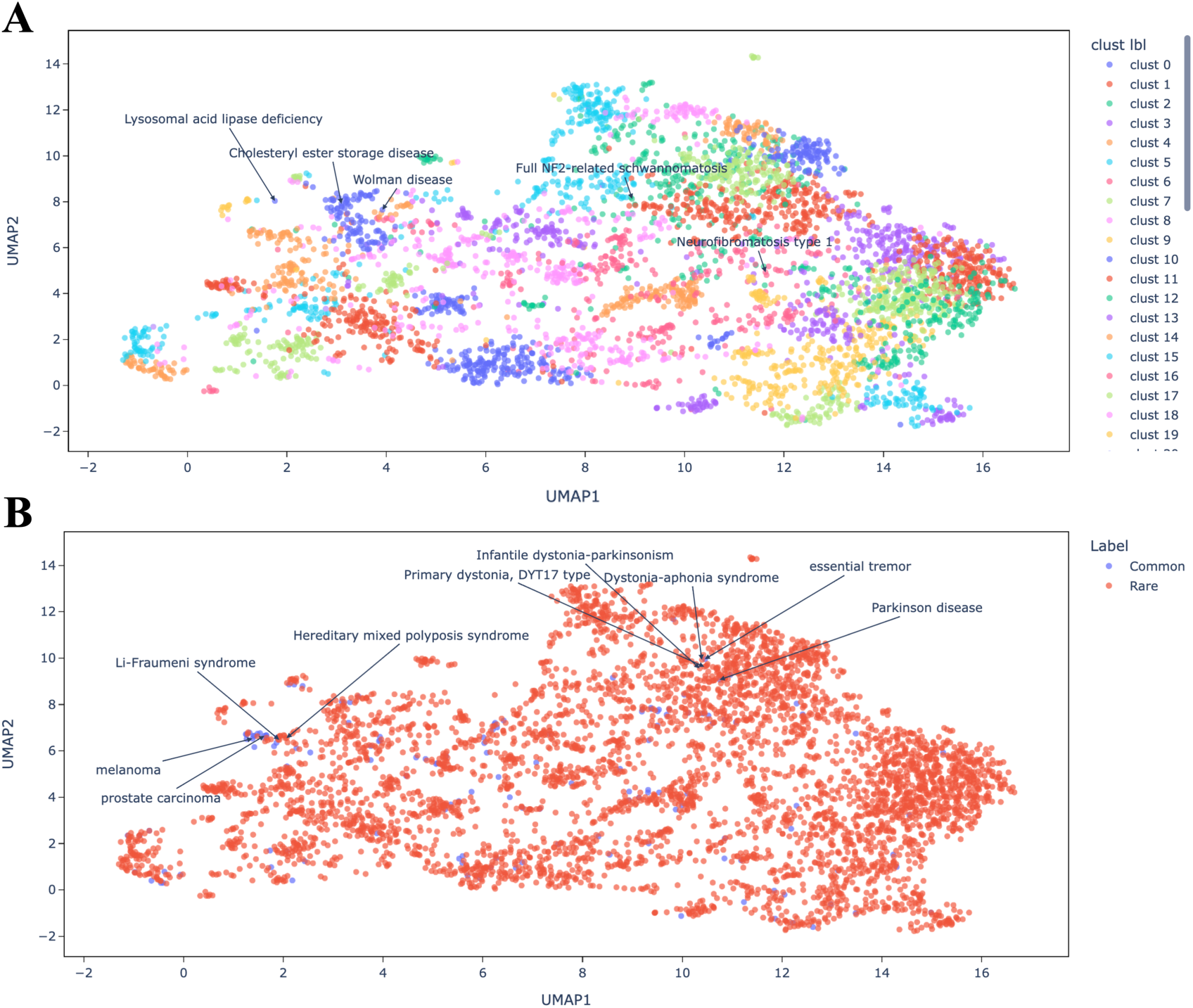
Dis2Vec Derived Embeddings Enables Disease-Maps and Unsupervised Clustering of Rare and Non-rare Disorders. Dis2Vec generates biologically grounded disease representations that support unsupervised clustering of both rare and non-rare diseases, producing a coherent disease map (see Methods). **(A)** Using UMAP to project the 128-dimensional Dis2Vec embeddings into two dimensions revealed strong concordance between UMAP visualization and the unsupervised Gaussian Mixture clusters. Cluster quality was evaluated using multiple positive and negative biological controls. For example, Lysosomal Acid Lipase Deficiency (LAL-D) and its subtypes-Cholesteryl Ester Storage Disease and Wolman Disease-were correctly grouped within Cluster 40, despite minor separations in UMAP space caused by differing Orphanet annotations. In contrast, Neurofibromatosis type 1 (NF1) and type 2 (NF2) were placed in distinct clusters, consistent with their distinct pathogenic mechanisms: NF1 is driven by Ras-signaling dysregulation, whereas NF2 involves loss of the Merlin protein, resulting in different clinical phenotypes. **(B)** The resulting disease map integrates rare and non-rare disorders according to shared biological mechanisms. Non-rare neurological diseases such as Parkinson’s disease and essential tremor cluster alongside rare hereditary dystonias and Parkinsonian syndromes, while non-rare cancers like melanoma and prostate carcinoma co-locate with rare hereditary cancer syndromes including Li-Fraumeni syndrome and Hereditary Mixed Polyposis Syndrome, illustrating the shared molecular foundations across the disease spectrum. This disease map also exhibits a rich, hierarchical organization. For instance, the cluster of immunodeficiencies (Cluster 0) is located near a cluster of immune-system cancers and autoimmune diseases (Cluster 31), which in turn is adjacent to clusters of autoimmune blood disorders (Clusters 5 and 47). This demonstrates a logical, higher-order structure between related disease groups. Crucially, the map seamlessly integrates rare and non-rare diseases based on shared biology (Figure 6B). Non-rare neurological disorders like Parkinson’s Disease are clustered with rare hereditary forms of Parkinson’s and other dystonias (Cluster 37), while non-rare cancers like Melanoma co-locate with rare hereditary cancer syndromes (Cluster 44), supporting our central hypothesis.

To further probe the accuracy of this disease map, we assessed its performance on specific positive and negative controls.

- **Positive Control:** Lysosomal Acid Lipase Deficiency (LAL-D) and its subtypes are all caused by mutations in the single gene LIPA. Dis2Vec correctly groups these disorders together in Cluster 40, overcoming differences in their clinical annotations in the source database.
- **Negative Control:** We tested two neurofibromatosis disorders, NF1 and NF2, which have similar names but are caused by distinct pathogenic mechanisms (Ras-signaling vs. Merlin protein loss). Dis2Vec correctly separates them into distinct and appropriate clusters to a RAS-related cancer cluster for NF1 and a neurological cancer cluster for NF2-demonstrating its ability to make fine-grained mechanistic distinctions.

### Explainability Contrast with a Knowledge Graph Foundation Model

The results suggest that agentic and graph foundation systems may benefit from an explicit disease-representation layer when evaluated on rare-disease cross-evidence generalization. Specifically, the limitations observed in agentic systems are consistent with the performance of state-of-the-art knowledge graph foundation models, such as TxGNN, when evaluated under the same strict cross-evidence generalization conditions (Figure 5A). To diagnose the cause of these limitations, controlling the training corpora and retraining the models is essential. This rigorous approach allows us to measure generalizability to entirely unseen data, thereby testing their capacity to mechanistically learn biology and provide meaningful predictions for questions falling out-of-distribution of the training corpus. To better understand these differences, controlled retraining allows graph models to be evaluated under held-out conditions that reduce topological leakage and test disease-level generalization. Consequently, in this section, we deeply examine the GraphML framework (Protocol 1). Unlike large language models, graph architectures afford us the distinct capability to systematically remove topological biases from the underlying networks prior to retraining. This enables us to strictly control for shortcut learning and isolate the model’s ability to extract underlying biological principles rather than overfit to training data. For example, while TxGNN performs exceptionally well on conventional within-graph tasks, it remains highly susceptible to the data-density biases present in clinical trial data, where only a small fraction of the total biological space has been empirically explored. This structural reliance ultimately leads to restricted generalizability and biologically ambiguous reasoning.

To complement the quantitative metrics presented in the Results section, we evaluated the relative capacities of TxGNN (original published model) and Dis2Vec to retrieve well-established mechanistic links between rare and non-rare diseases. To do so, we selected three classic disease pairs from the literature:

1. **Marfan Syndrome and Thoracic Aortic Aneurysm (TAA):** Mechanistic insights from Marfan syndrome, caused by FBN1 mutations, inform the role of extracellular matrix defects in TAA [55].
2. **Maturity-Onset Diabetes of the Young (MODY) and Type 2 Diabetes:** MODY, a rare monogenic form of diabetes, has clarified key pathways in beta-cell dysfunction relevant to the non-rare Type 2 Diabetes [56].
3. **Gaucher Disease and Parkinson’s Disease:** Gaucher disease, a lysosomal storage disorder caused by GBA mutations, provides a clear mechanistic link to Parkinson’s, as carriers of a single GBA mutation have a significantly higher risk [57].

Even though this specific analysis evaluated the published version of TxGNN without removing drug nodes or controlling for historical clinical trial biases, TxGNN Explorer [58], the platform’s native explainability module, nevertheless highlighted cases where the retrieved paths and therapeutic suggestions were less aligned with the expected disease mechanisms (Figure 7B). These examples suggest that graph foundation models may require additional disease-level grounding to support reliable rare-to-non-rare mechanistic transfer. TxGNN Explorer successfully connected MODY to Type 2 diabetes but failed on the remaining two disease pairs, generating biologically implausible pathways. Specifically, it linked Marfan syndrome to schizophrenia and Gaucher disease to entirely unrelated symptoms. Furthermore, the top therapeutic candidates recommended by TxGNN in these failed cases lacked clinical relevance to the target indications. For Marfan syndrome, the model recommended imatinib (Gleevec), a small-molecule tyrosine kinase inhibitor that targets the BCR-ABL fusion protein driving oncogenic conditions like chronic myeloid leukemia. For Gaucher disease, it recommended nusinersen, an antisense oligonucleotide designed to compensate for the loss of survival motor neuron protein in spinal muscular atrophy. This inability to recover literature-supported pathways underscores a fundamental limitation in TxGNN, demonstrating that it struggles to maintain biological fidelity when forced to make inferences outside its explicit training topology.

**Figure 7:**
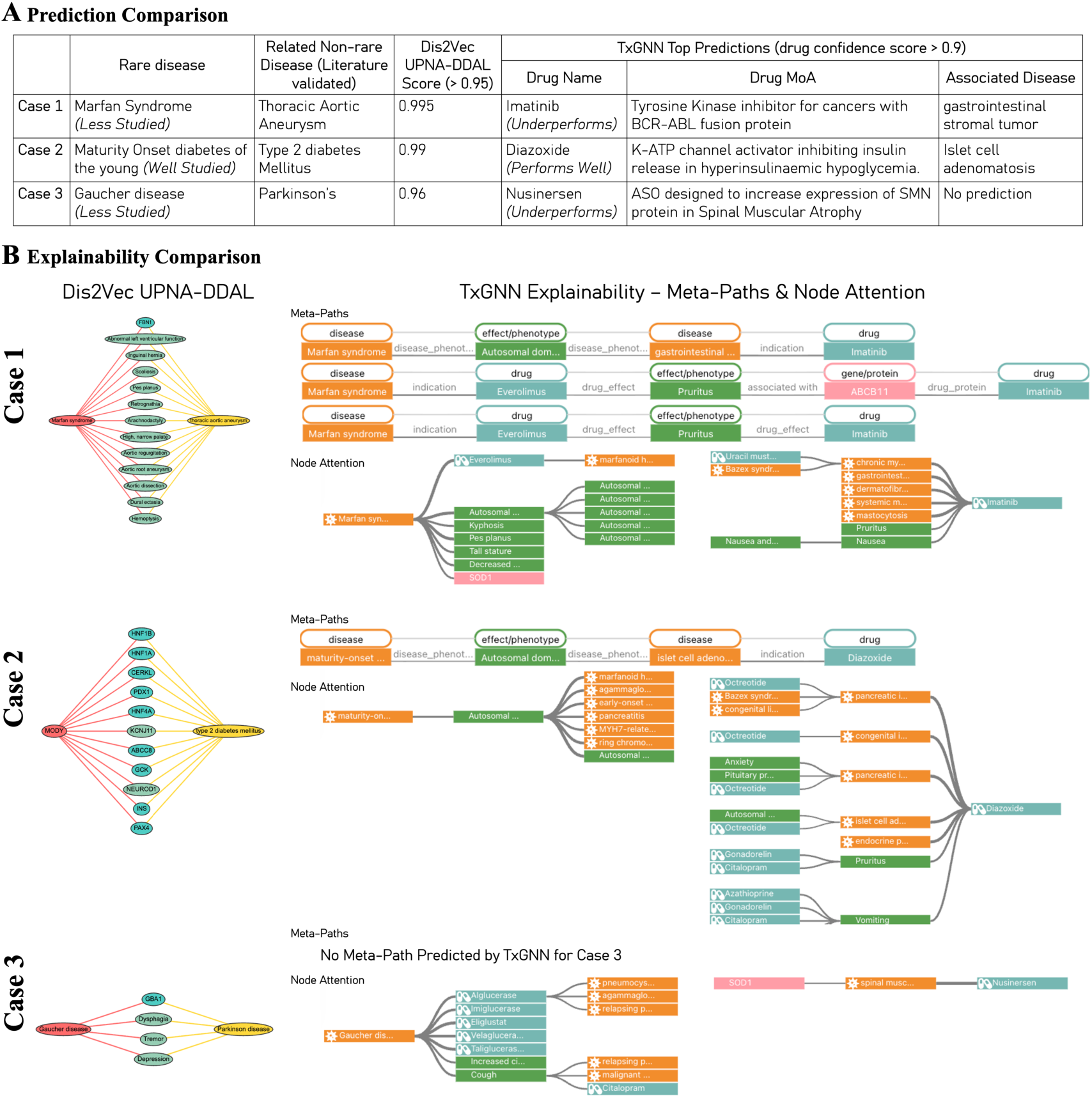
Literature validated rare and non-rare diseases benchmark on Dis2Vec UPNA-DDAL and TxGNN. **(A) Prediction Comparison:** TxGNN underperforms in Case 1 and Case 3, which involve sparsely studied rare diseases, but performs well for well-studied rare diseases like Maturity Onset Diabetes of the Young in Case 2. In Case 1, Marfan syndrome reveals how extracellular matrix defects contribute to non-rare aortic aneurysms and arterial disease [55]. In Case 3, Gaucher disease-a lysosomal storage disorder from biallelic GBA mutations-shows that heterozygous GBA variants, though rare, markedly increase Parkinson’s disease risk, linking lysosomal dysfunction to neurodegeneration [57]. **(B) Explainability Comparison:** We present the sub-networks contributing to Dis2Vec UPNA-DDAL and TxGNN. Dis2Vec provides pathway-level reasoning linking relevant bio-entities to the rare and non-rare diseases in Case 1 and Case 2, whereas TxGNN’s predictions are less specific. For example, in Case 1, TxGNN’s less relevant prediction arises from prioritizing broad phenotypes-such as autosomal dominant inheritance-that lack specificity to Marfan syndrome. Lastly, although TxGNN performs well in Case 2, the meta-paths and node attentions connect MODY and Type 2 diabetes in a relatively nonspecific manner within the broader context of diabetes, since both are subtypes of Diabetes mellitus and involve pancreatic or insulin-related dysfunction.

In contrast, Dis2Vec framework addresses part of this gap by encoding disease etiology and phenotype into a dedicated disease-level representation space by capturing the underlying disease etiology rather than memorizing direct graph shortcuts. Dis2Vec assigned high association scores to all three biologically linked pairs (Figure 7A) by extracting higher-order relationships from the MechDist network, which is optimized for the DDAL task. As illustrated in Figure 2B, these robust predictions are driven by higher-order information traced back to the underlying disease–gene and disease–symptom associations used to construct MechDist. By embedding this multi-modal biological layer, Dis2Vec delivers interpretable and mechanistically grounded predictions that remain robust even across challenging, cross-evidence validations.

## Discussion and Conclusion

The AI revolution in drug discovery has dramatically shifted paradigms. The fundamental question is no longer whether larger architectures can memorize broader sheets of historical associations, but whether they can capture transferable biological mechanisms that support genuine therapeutic transferability [59, 60]. Traditional evaluation strategies, such as random train-test-validation splits, frequently rely on label-driven or within-graph metrics. Consequently, they have become insufficient to distinguish superficial pattern matching from true mechanistic understanding [61]. Aligned with this shift, our work demonstrates the necessity of engineering mechanism-centered graphs rather than relying on raw knowledge graphs to disentangle structural network biases and optimize the global signal-to-noise ratio [35].

Another fundamental shift centers on how translationally relevant science is conducted. Traditional hypothesis-driven research typically begins with a proposed mechanism or model and subsequently seeks supporting evidence. However, when applied to large-scale foundation models, this sequence is increasingly prone to overfitting, confirmation bias, and superficial pattern matching. In contrast, the most reliable path to genuine progress now requires a benchmark-first philosophy, beginning with the intentional design of rigorous, orthogonal evaluation frameworks that independently test for true mechanistic understanding before any model is developed [62, 63]. The IxIDN and ORDON benchmark pair [40] exemplifies this philosophy in the DDAL task, providing a highly rigorous, decision-oriented testbed. Its orthogonal architecture anchors evaluation in real-world clinical trial investment decisions while utilizing stringent, biology-aware negative controls derived from ORDO. By ensuring minimal overlap between training and testing modalities, this setup provides a principled test of cross-evidence generalization rather than label memorization or information leakage, directly addressing a critical bottleneck in building trustworthy translational AI. Ultimately, our results demonstrate that even state-of-the-art LLMs and agentic AI systems are powerful for biomedical synthesis and tool orchestration, but this benchmark suggests they still need structured, biologically grounded disease representations to support reliable mechanism transfer in rare-disease settings. In contrast, mechanism-centered representations such as Dis2Vec exhibit superior generalization under strict inductive constraints. This divergence highlights the dual value of constructing mechanistic-centered graphs over raw knowledge graphs, and pairing this architecture with orthogonal, biology-grounded benchmarks to effectively guide the next generation of biomedical AI.

### A Continuous Mechanistic Landscape of Human Disease

A central insight from this work is the value of moving beyond discrete, label-driven disease definitions toward a **continuous, composite representation** grounded in shared biological mechanisms. Rather than treating diseases as isolated ontological categories, Dis2Vec embeds them as points in a continuous vector space whose geometry reflects underlying pathophysiology. The barycentric compositionality observed in the embeddings (see Supplementary Section on Compositionality) provides direct evidence that this continuous view is biologically meaningful: rare and non-rare disorders can be positioned along the same mechanistic gradients, allowing knowledge to transfer smoothly across what are traditionally viewed as separate clinical entities.

This continuous definition manifests most clearly in the unsupervised disease map. Dis2Vec organizes the entire landscape into coherent “continents” of related biology, such as autoimmune, neurodegenerative, or metabolic clusters, that are arranged according to higher-order mechanistic programs rather than clinical specialty or superficial phenotypic similarity. Within these continents, rare and non-rare disorders are seamlessly integrated around shared molecular and phenotypic mechanisms. For example, Parkinson’s disease co-clusters with its rare hereditary variants and related dystonias, demonstrating that many rare conditions are not isolated outliers but extreme points on the same mechanistic spectrum as more non-rare disorders.

By revealing this higher-order organization, Dis2Vec offers a practical framework for improving the efficiency of translational science. It enables the identification of therapeutic “baskets” [29] that span rare and non-rare diseases sharing core pathway dysregulation, thereby addressing a long-standing data-scarcity barrier in rare-disease research. Ultimately, these findings support the hypothesis that decision-oriented, mechanism-centered benchmarks can drive more reliable and interpretable disease representations than those derived from conventional label-driven or same-graph approaches.

### Why Agentic and Graph Models Need Disease-Level Grounding

A dedicated evaluation under strict cross-evidence generalization revealed clear and category-specific limitations. Compared to our Dis2Vec-UPNA-DDAL, both LLMs and agentic AI systems showed less discrimination between real therapeutic transfer pairs and mechanistically distant negatives, suggesting that retrieval and tool use alone may not be sufficient when rare-disease evidence is sparse or fragmented. This finding is aligned with the observed gap that LLMs do not reliably learn or reason over graph structure when graphs are presented as text [64]. Surprisingly, agentic systems equipped with external tools and retrieval performed no better — and in some cases slightly worse — than their underlying LLMs. This is likely because the benchmark is heavily enriched in rare diseases, where structured cohort data and well-curated pathway resources are scarce; tool-augmented retrieval therefore returns little actionable information, while LLMs can still fall back on memorized literature.

Knowledge-graph and GraphML models exhibited a different but equally fundamental weakness. For instance, TxGNN, the state-of-the-art drug repurposing model, exhibited substantial performance degradation (AUROC = 0.575; Figure 5A) once drug nodes and benchmark edges were removed from training, revealing heavy reliance on shortcut learning rather than mechanistic inference. To probe whether this failure is architectural rather than a consequence of PrimeKG’s heterogeneity, we modified TxGNN’s meta-paths to operate on the homogeneous, unipartite MechDist graph — the same graph used to train Dis2Vec and the GCN baseline. Surprisingly, TxGNN performed even worse on MechDist (AUROC = 0.449, below random; Figure 5C), demonstrating that TxGNN’s relational meta-path inductive bias is misaligned with homogeneous disease–disease graphs and cannot recover therapeutic transferability even when provided a purpose-built mechanistic graph. In contrast, the simple GCN trained on MechDist achieved substantially better discrimination (AUROC = 0.780; Figure 5D), confirming that the bottleneck is the architecture, not the underlying graph. These findings indicate that current graph foundation models are not yet effective at capturing the higher-order network structures — multi-hop pathways and hierarchical biological organization — that are essential for reliable therapeutic transferability. Similar observations have been reported across a broad range of KG-based drug-disease association learning methods, from heterogeneous graph transformers (e.g., HGTDR) to other state-of-the-art approaches such as DeepDR and HeTDR, further reinforcing that shortcut learning and limited generalization remain pervasive challenges in this class of models.

Collectively, these findings support a complementary role for Dis2Vec in providing a benchmarked disease-representation layer that can augment foundation models and agentic systems with structured mechanistic context, paving the way to capture higher-order mechanistic information critical for genuine therapeutic transferability. Rather than replacing agentic AI systems, Dis2Vec framework provides a biologically grounded disease-representation and benchmarking layer that can make agentic biomedical reasoning more reliable in rare-disease indication expansion.

### Limitations and Future Directions

Although Dis2Vec is an important first step, several limitations and future directions remain. First, some notable associations remain difficult to explain. For example, RA and IBM (Inclusion Body Myositis) has a strikingly lower association score (0.34) than the other IIM disorders, whereas some less well-known autoimmune disorders, such as Autoerythrocyte Sensitization Syndrome, showed a higher association probability than all IIMs (0.97). Such cases may present interesting directions for research on shared disease mechanisms and suggest novel repurposing opportunities.

Second, Dis2Vec UPNA-DDAL does not identify associations for therapeutics that target a specific molecular lesion rather than a broader biological pathway or phenotype. Dis2Vec UPNA-DDAL predicted only 21 of the 55 indications for Ataluren, a drug that works by allowing read-through of nonsense mutations, because many of these diseases are linked by a shared molecular mechanism rather than a shared pathophysiology. In contrast, therapies that target a conserved pathophysiological mechanism, such as setmelanotide, were strongly supported by the model, reinforcing the validity of its biological-first approach.

Third, while the global structure of the Dis2Vec map is biologically coherent, we also identified limitations. The placement of individual diseases can be sensitive to variations in their phenotypic annotations. For example, some subtypes of NF1, despite being linked to the same causal gene, appear in a different cluster from the primary NF1 entry. This suggests that while Dis2Vec produces a robust global disease architecture, its local precision can be influenced by annotation inconsistencies in the source data, highlighting an area for future refinement.

Finally, several promising directions remain for future work. An important next step is to extend Dis2Vec toward a JEPA-like biomedical architecture [65], in which disease context, molecular evidence, phenotypic profiles, patient-level observations, and therapeutic perturbations are encoded into a shared latent space. In such a framework, models could be trained to predict masked or future biological states directly in embedding space, moving indication expansion from retrospective association learning toward predictive disease-world modeling for translational AI. Patient-level data, including symptoms and genomic sequences, could also be mapped onto the Dis2Vec space to support diagnosis and patient stratification. In addition, comorbidity patterns, real-world evidence, and multi-omics data could be incorporated to further enrich the representation space.

## Methods

### Data Preparations

#### Indication Expansion Investment Decision Network (IxIDN) as Positive Control

The IxIDN is created as described in [40]. This network serves as our primary reference dataset for validating the DDAL task, since it recapitulates drug repurposing investment decisions made in the pharmaceutical industry.

#### Orphanet Rare Disease Ontology Negative-network (ORDON) as Negative Control

To quantify the rate of false negatives, we used the ORDON [40], this disparity network is created based on the maximization of the LCA distance in the Orphanet Rare Disease Ontology (ORDO) [42]. The ORDON pairs, although sourced from the rare-disease-focused ORDO, span multiple therapeutic areas and were chosen to enforce the strongest possible mechanistic boundaries in the benchmark.

#### Selection of Gene Associations with Strong Genetic Evidence

Because Dis2Vec was designed to support representation learning across both rare and non-rare diseases, we used the OpenTargets platform as the primary source of disease–gene associations. OpenTargets integrates evidence from multiple resources, including Orphanet, while also capturing genetic associations for non-rare diseases through sources such as GWAS and literature-derived evidence.

To reduce noise, we used the OpenTargets Overall Score and retained only disease–gene links supported by at least 10 independent pieces of evidence, including both GWAS-derived and NLP-based association evidence. After applying this filter, we obtained a total of 6,654 diseases. As shown in Figure S2, 47% of diseases had both disease–gene associations and curated symptom annotations, whereas 53% were missing one of these two evidence types. This level of missingness highlights the importance of representation learning approaches such as Dis2Vec, which can leverage available heterogeneous evidence across the rare–non-rare disease spectrum.

#### MechDist Network

To assess whether the disease representations learned by Dis2Vec is able to accurately predict whether a pair of diseases will be evaluated for response to the same drug (Investment Decision) or whether they would fall into our class of ORDO ontological negatives (ORDON). We constructed this task as follows. We have a disease association graph *G*. *V* is the set of disease nodes. The graph has 4 different types of edges:

- *E*_1_: Edges based on Gene Ontology (GO), derived based on minimizing LCA.
- *E*_2_: Edges derived based on (Human Phenotype Ontology) HPO.
- *E*_3_: Edges from IxIDN [40].
- *E*_4_: Negative edges from ORDON [40].

We defined ontological similarities by minimizing LCA distances within the hierarchical structures of the GO Biological Processes and the HPO [66], combined with anchoring disease-gen and -symptom associations to these hierarchies (see Figure S3 and Figure 2). MechDist was constructed by computing LCA distances for every disease pair across these two modalities. We applied a threshold of 0.2 to filter pairs with high distances (Figure S5) to obtain *E*_1_ and *E*_2_. The reason for integrating a rank aggregation method [67, 68] is to address the substantial data incompleteness in rare and non-rare diseases, 53% of diseases lack either genetic associations or curated symptoms (see Figure S2). Hence, *G_MechDist_* = (*V*, *CRank*(*E*_1_, *E*_2_) − *E*_3_) forms the MechDist network, which we use for training UPNA-DDAL, a disease association prediction decoder on Dis2Vec embeddings. This process filtered out 1,944 diseases from the initial pool of 6,654; these diseases did not meet the quality threshold required to accurately repurpose knowledge and assign reliable Dis2Vec representations. Consequently, the current Dis2Vec embeddings cover 4,710 diseases (4,565 rare diseases and 145 non-rare diseases).

#### Dis2Vec Embeddings

We used Node2Vec to translate the topology of the MechDist graph into disease-level embeddings, which we refer to as Dis2Vec embeddings. Each disease was represented by a 128-dimensional vector. We selected an embedding dimension of 128 because it provided the best performance in downstream tasks. Node2Vec was trained with a learning rate of 10^−5^, *β*_1_ = 0.9, and *β*_2_ = 0.999. For *G*_MechDist_, we used a walk length of 10, 100 walks per node, a window size of 10, and return and in-out parameters of *p* = 1 and *q* = 1, respectively.

#### Cross-Evidence Generalization Setup to Evaluate Knowledge Graph Based Approaches

Cross-evidence generalization for KG-based models follows Protocol 1 described above. Concretely, all IxIDN edges (*E*_3_) and ORDON edges (*E*_4_) are removed from the training graph, and all drug nodes are removed from the KG to prevent shortcut learning via memorized drug-disease paths. The model is then trained exclusively on the remaining MechDist graph with *E*_3_ and *E*_4_ held-out. For KG-based approaches, like TxGNN, drug nodes are also excluded.

The UPNA-DDAL (disease-disease association decoder) is trained using Dis2Vec embeddings learned from MechDist. Cross-evidence generalization is evaluated by predicting the investment decision and ORDON links (Figure 5). Since the network properties of MechDist, IxIDN, and ORDON are significantly different (see Figure S5B and Figure S6), the cross-evidence generalization task ensures model stability under distribution shifts [69] and the ability to bypass topological shortcuts [33, 70, 71].

#### Dis2Vec-UPNA-DDAL Model Architecture for Disease Association Learning

Similar to the two-shot learning approach proposed by Chatterjee et al. [38], we feed the Dis2Vec embeddings into a three-layer multi-layer perceptron (MLP) decoder [72, 73]. The decoder is trained and validated in a transductive setting on the MechDist network for the DDAL task. To train Dis2Vec-UPNA-DDAL, we construct a cross-evidence generalization task for DDA prediction, formulated as link prediction. Specifically, we remove the investment-decision edges, *E*_3_, from *G*_MechDist_ because these links are reserved for evaluating cross-evidence generalization, a form of inductive testing. Thus, training and validation of the Dis2Vec-UPNA-DDAL model are performed on the disease graph *G*_train_ = *G*_MechDist_, while cross-evidence generalization is evaluated on the test graph *G*_test_ = *G*_ID_ = (*V*, *E*_3_). During training on *G*_train_, we use random negative sampling from *G*_train_. In contrast, for evaluation, we use the ORDON edges, *E*_4_, from *G*_ORDO_ = (*V*, *E*_4_) as a negative-control set to assess the model’s ability to reject unsupported investment-decision links.

#### Unsupervised Clustering of Diseases

We used the Gaussian Mixture model from sklearn directly on disease vectors (obtained from Node2Vec) to identify 50 clusters. The number of clusters is empirically obtained by assessing positive and negative controls such as LALD (Lysosomal acid lipase deficiency) and CESD (Cholesteryl ester storage disease) as a positive control, and NF1 (Neurofibromatosis type 1) and NF2 (Full NF2-related schwannomatosis) as a negative control. Next, we used UMAP to reduce dimension and visualize the clusters. We observe that the reduced dimensions by UMAP largely agrees with the clusters obtained directly from the 128 dimensions of Dis2Vec.

#### Evaluation of Dis2Vec Across GraphML, LLM, and Agentic AI System Categories

##### Cross-Evidence Generalization: Four Evaluation Protocols

The term *cross-evidence generalization* refers to evaluating a model on data that is entirely independent of its training signal. Because KG-based models, LLMs, agentic AI systems, and disease-drug ranking models have fundamentally different architectures and constraints, four distinct protocols were applied:

##### Protocol 1 — GraphML on homogeneous/heterogeneous graphs (unipartite graph vs. KG; Dis2Vec/UPNA-DDAL, TxGNN)

Cross-evidence generalization is implemented as a strict inductive cross-evidence generalization test: (i) all IxIDN edges (*E*_3_) and ORDON edges (*E*_4_) are removed from the training graph; (ii) all drug nodes are removed from TxGNN’s heterogeneous KG (PrimeKG) to prevent shortcut learning via memorized drug-disease paths — MechDist contains no drug nodes; (iii) the model is retrained on the remaining graph; (iv) performance is evaluated exclusively on the held-out IxIDN positives and ORDON negatives as out-of-training edges. This ensures no therapeutic association signal of any kind leaks into training and is the most stringent of the four protocols.

##### Protocol 2a — LLMs / parametric scoring (GPT-5.5, Claude-Opus-4.7, Gemini-3)

At present, it is difficult to retrain LLM models in a way that reliably audits whether drug-trial knowledge has been removed, altered, or selectively stripped from the training corpus. Accordingly, Protocols 2a and 2b should be interpreted as practical decision-support baselines rather than fully controlled architectural comparisons. Their purpose is to test whether current general-purpose and tool-augmented systems can recover the benchmark signal without a dedicated disease-representation layer. Cross-evidence generalization is tested via structured prompting with no tool augmentation: each model scores disease pairs from its internal parametric knowledge alone. Each model was provided the blinded IxIDN and ORDON benchmarks released in [40]. No tool calls were made.

##### Protocol 2b — Agentic AI systems / tool-augmented scoring (Phylo, Edison Scientific)

Agentic systems extend the LLM core with tool use, retrieval, and multi-step reasoning. Cross-evidence generalization is tested using the **same structured prompt as Protocol 2a**, but the model may additionally call external tools or databases (e.g., MCP calls, biomedical knowledge bases) at inference time before assigning a DDAL score. Evaluation is identical: AUROC, AUPRC, and score separation (Δ) against IxIDN/ORDON labels. Phylo and Edison Scientific were accessed via their web interface. This constitutes a weaker but practically realistic form of cross-evidence generalization for Protocols 2a and 2b, as both system types may have encountered drug or trial information during pretraining.

The following is the exact prompt submitted verbatim to all Protocol 2a and 2b systems, together with the blinded benchmark CSV (IxIDN_dis2vec_filtered_benchmark.csv):

*“You are a disease-disease association learning (DDA) agent. Your task is to calculate DDA(d1, d2), which returns an association score between 0 and 1 for two diseases to quantify the extent these diseases share mechanisms of action. Strictly, do not use clinical trials and drug datasets to implement the DDA function*.

*For each disease pair in the attached csv file, compute DDA and add a new column to the csv file named DDA to export the DDA scores for each row. Make sure in the final csv file you are producing exact same columns as attached without any modification and just add a single new column named DDA.”*

## Data Availability

All data used in this study were obtained from public sources, including Orphanet, OpenTargets, the Gene Ontology, and the Human Phenotype Ontology. The Dis2Vec embeddings, all UPNA-DDAL predictions, and the GO/HPO term–term distance matrices derived from (LCA) calculations are available at https://zenodo.org/records/21251215.

## Code Availability

The codes that support the findings of this study are openly available on our GitHub at https://github.com/alxndgb/Dis2Vec.

## Acknowledgements

The authors acknowledge AstraZeneca’s Scientific Computing Platform (SCP) for providing the computational infrastructure and resources that enabled this research.

## Author Contributions

C.B.R., W.R.M., A.C., and P.R. conceived and designed the project. C.B.R., W.R.M., and A.C. wrote the manuscript and conducted data curation and preparation. C.B.R. conducted the data engineering and integration to produce MechDist and performed GraphML modeling and analysis. W.R.M. performed an in-depth evaluation of clusters and the Investment Decision benchmark. A.C. designed the machine learning architecture and tuned the hyperparameters. F.K. provided clinical insights and contributed to manuscript writing. P.H. performed biological validations and contributed to manuscript writing. M.U. and I.B. provided insights on knowledge graph engineering and model design and contributed to manuscript writing. W.D. contributed to evaluation of results. G.D.A. contributed to manuscript writing. T.D. designed the compositionality analysis of the Dis2Vec space and contributed to manuscript writing. J.C.M. performed an in-depth evaluation of Dis2Vec and TxGNN, designed the explainability contrast with a knowledge graph foundation model, and contributed to manuscript writing. S.L. offered clinical feedback on positive and negative controls and contributed to manuscript writing. S.C. interpreted the results, offered biological insights, and contributed to manuscript writing. T.E.R. provided guidance on designing the machine learning experiments and writing the manuscript. G.R. contributed to the design and strengthening of the IxIDN and ORDON benchmarks, supported the maturation and execution of the Dis2Vec codebase, played a key role in the validation, interpretation, and evaluation of benchmarks on LLMs and foundation models, and made significant contributions to manuscript writing. P.R. conceived the study design and contributed to manuscript writing.

## Declaration of generative AI and AI-assisted technologies in the manuscript preparation process

During the preparation of this manuscript, the authors used ChatGPT and Claude to assist with manuscript revision. The authors reviewed and edited the output as needed and take full responsibility for the content of the published article.

## Supplementary Information

### S1. Prevalence of Rare Diseases

**Figure S1:**
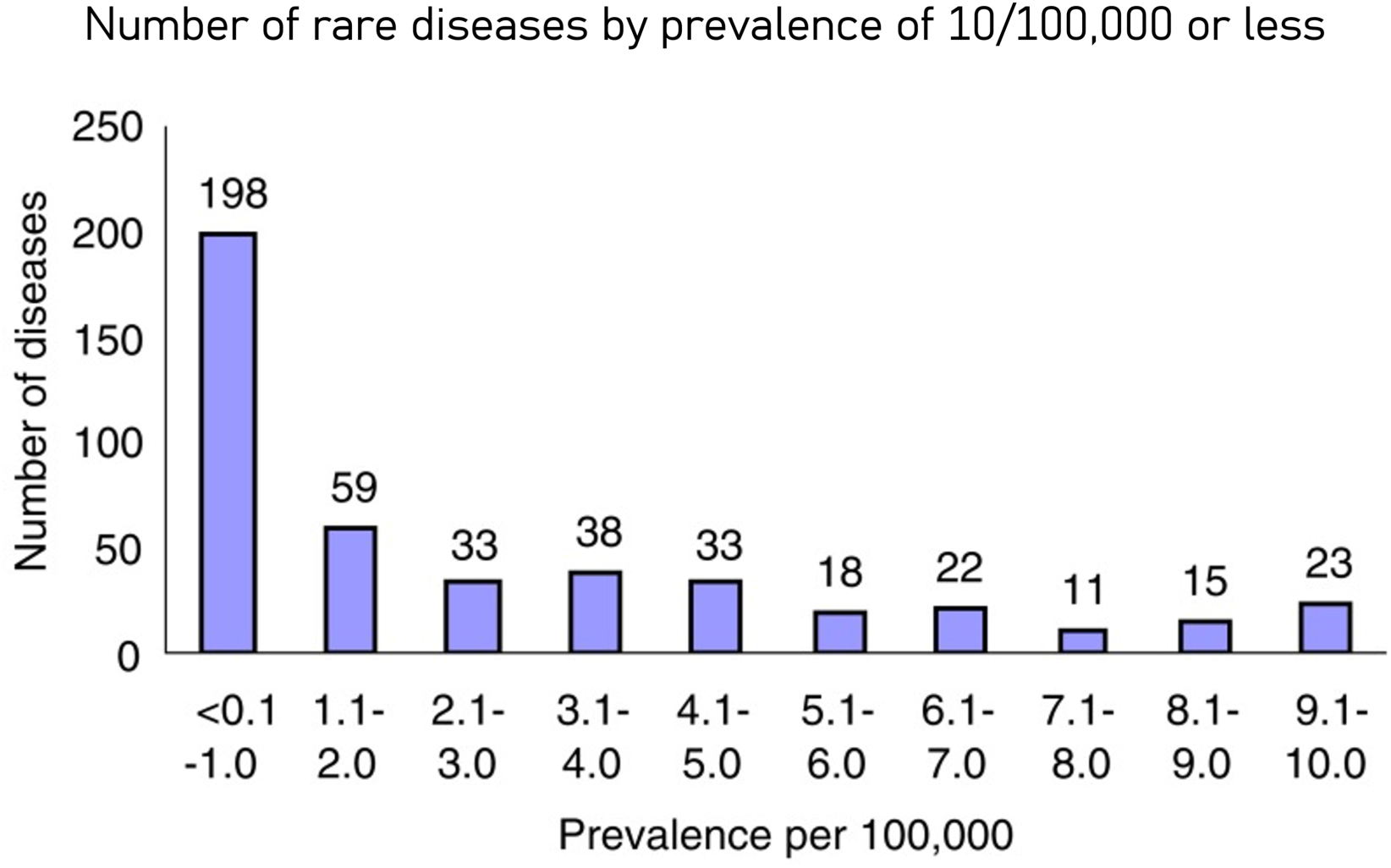
Prevalence of Rare Diseases. The distribution of prevalence of rare diseases follow a long tail, where majority of rare disease are ultra-rare, hence various phenotype are observed in the low prevalence rare diseases. Only a few rare diseases have comparably higher prevalence, hence are more studied. Lastly, rare diseases are heavily under diagnosed, making it difficult to obtain accurate prevalence of majority rare diseases. Source of the distribution plot: Courtesy of Boat and Field [74].

### S2. OpenTargets Disease Gene Associations

**Figure S2:**
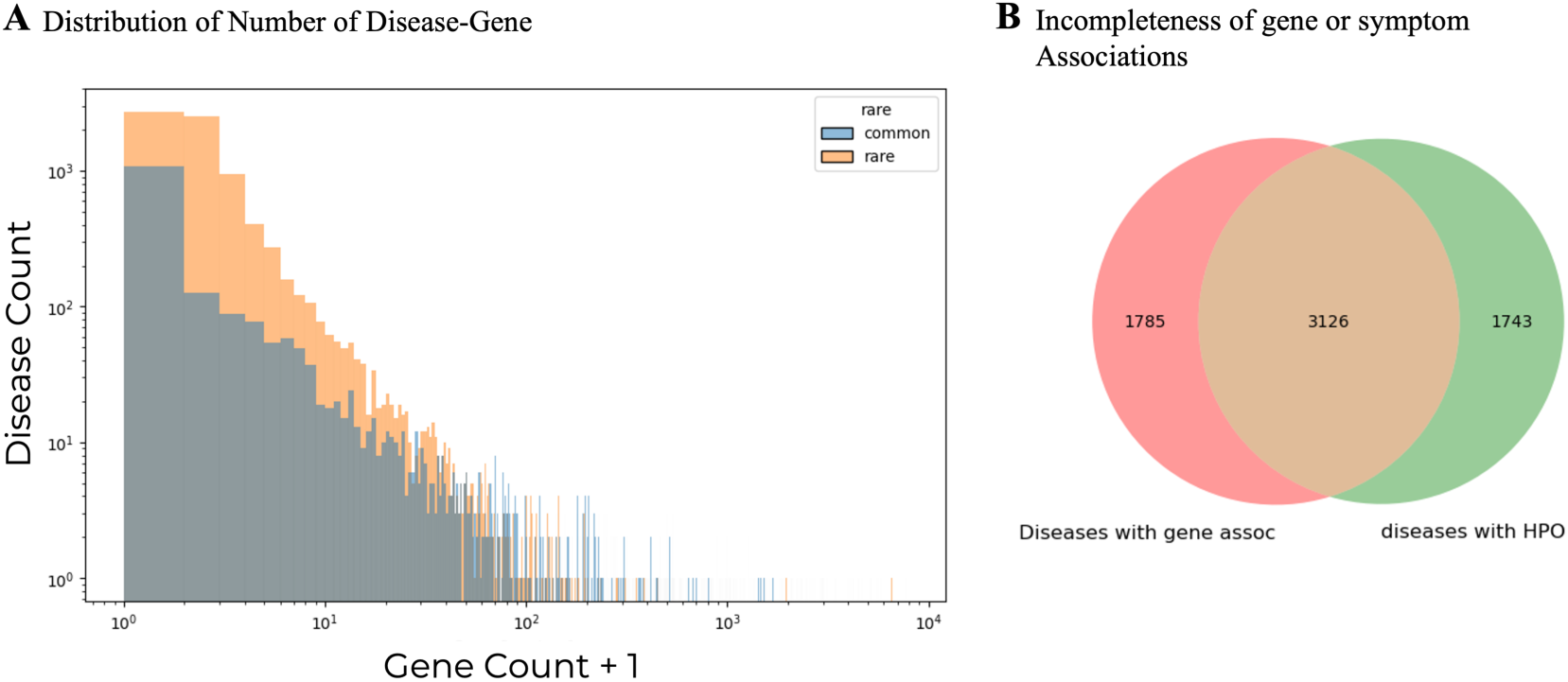
Extent of data Incompleteness for Both Rare and Non-Rare Diseases a Picture from OpenTargets Data. **(A)** Distribution of Number of Disease-Gene Associations from OpenTargets for non-rare and Rare Diseases. No filters are applied, for example diseases without genetic score are included if they have disease-gene associations according to *OpenTargets Overall Score*, which includes extended sources such NLP-based disease-gene associations. **(B)** Not all diseases have both symptom and gene associations, hence it is necessary to combine both modalities to obtain a comprehensive disease representation. This analysis shows that 47% (3,126 / 6,654) of diseases have both disease-gene associations and curated symptoms, demonstrating that 53% of diseases ([1,785 + 1,743] / 6,654) are missing one of the two evidence types (either disease-gene associations or curated symptoms).

### S3. Disease Similarity from Hierarchical Structure of Ontologies

**Figure S3:**
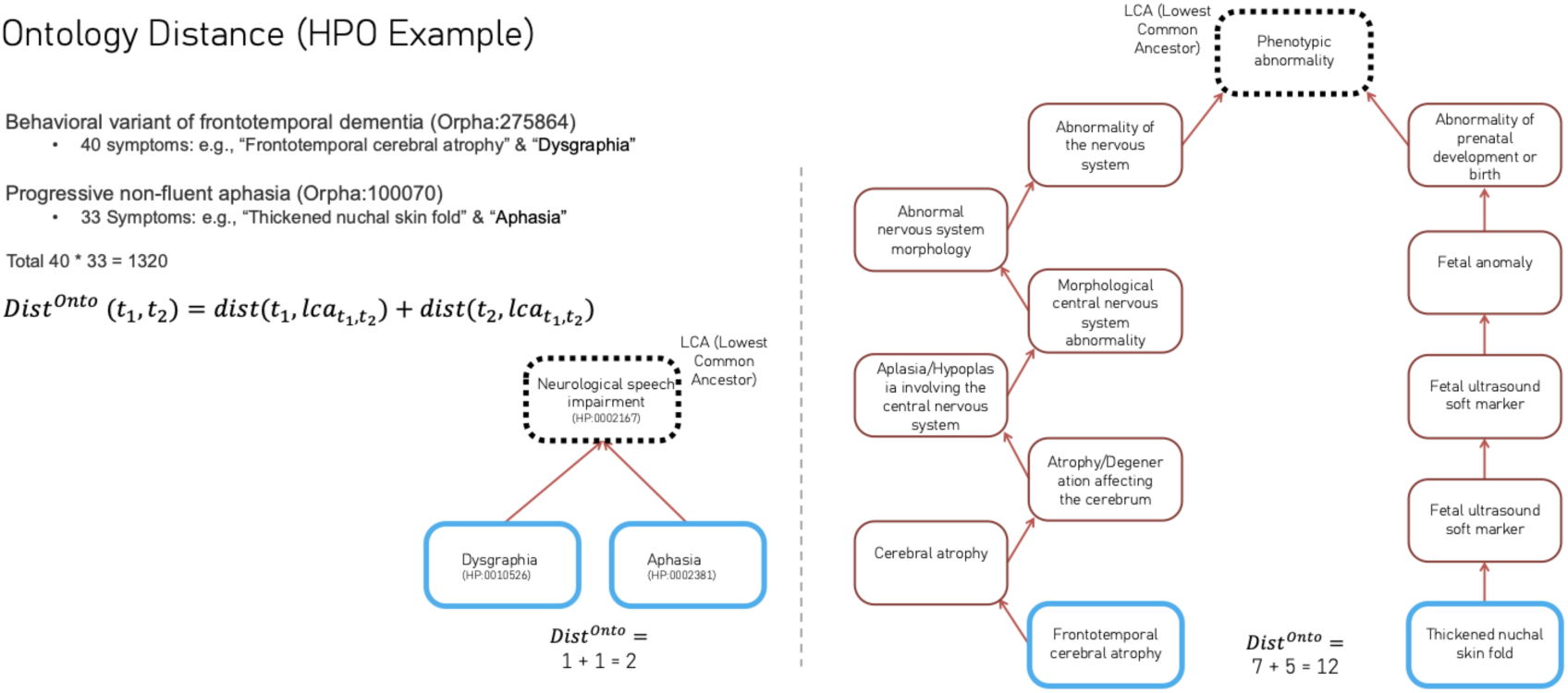
Distance to Lowest Common Ancestor.

**Figure S4:**
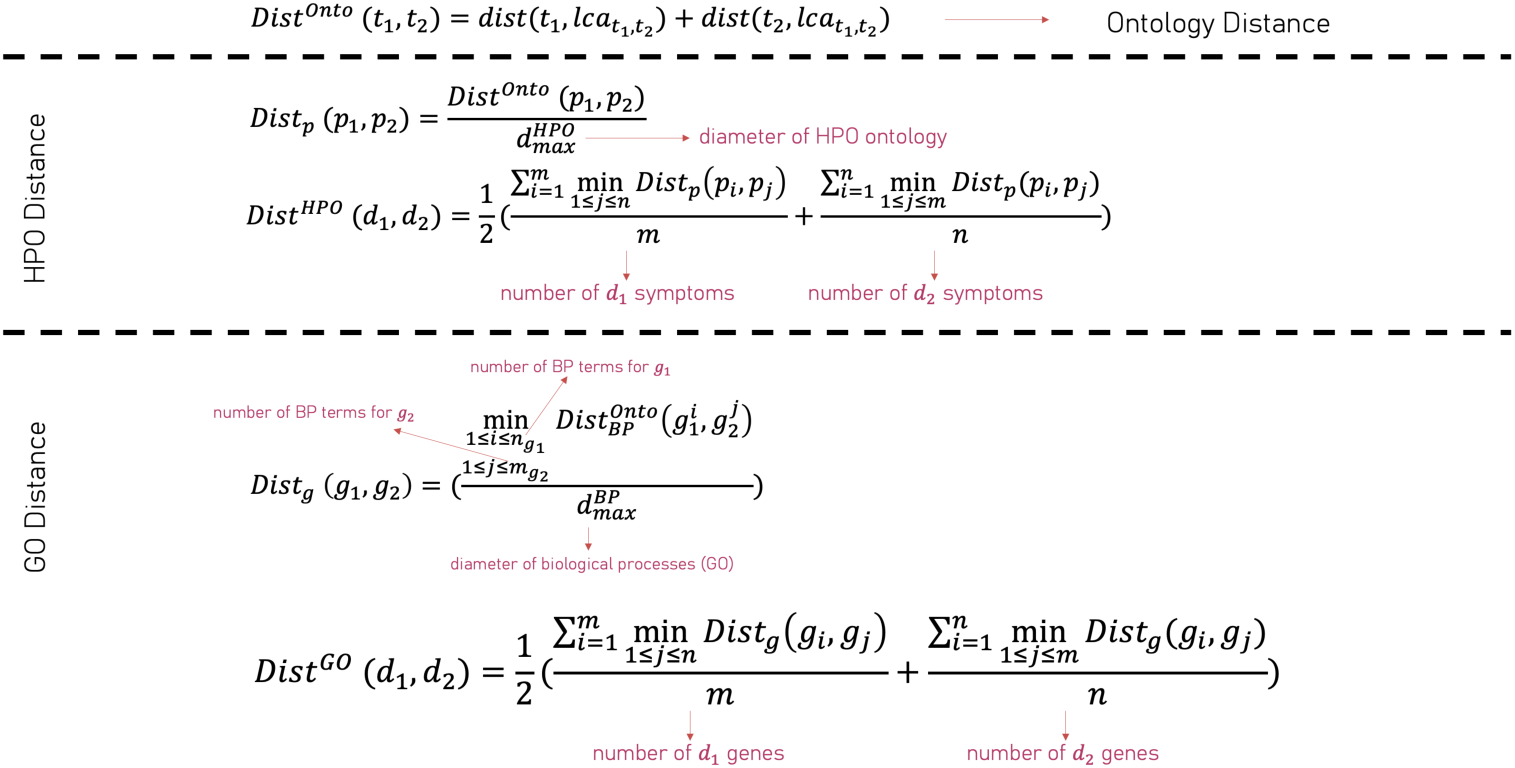
Disease Similarity by Minimizing LCA Distances. The diameter of a network is obtained by finding the longest shortest path in the network.

**Figure S5:**
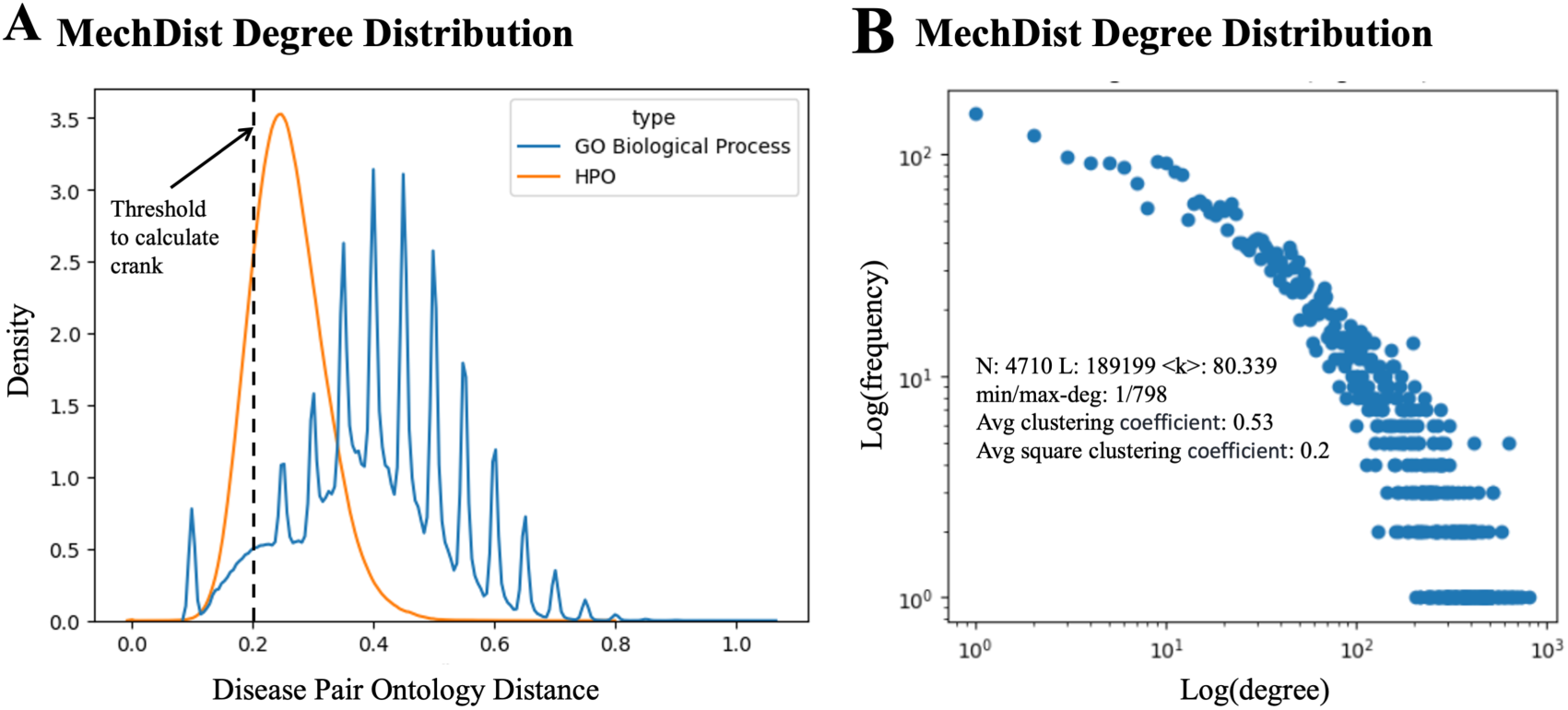
Disease-Pair Similarity Distance and MechDist Degree Distributions. (A) Distributions of disease-pair similarity distances computed by minimizing Lowest Common Ancestor (LCA) distances within the Gene Ontology (GO) Biological Processes and Human Phenotype Ontology (HPO) hierarchies, followed by harmonic mean projection to the disease level. The GO Biological Processes hierarchy exhibits clear oscillation; a similarity threshold of 0.2 was therefore selected to define disease–disease associations. **(B)** Degree distributions of the MechDist network. MechDist exhibits markedly different network properties and interaction patterns compared to the IxIDN and ORDON benchmarks (see Figure S6), making the cross-evidence generalization task from Dis2Vec-UPNA-DDAL to IxIDN and ORDON particularly challenging.

### S4. Degree Distribution of IxIDN and ORDON Benchmarks

**Figure S6:**
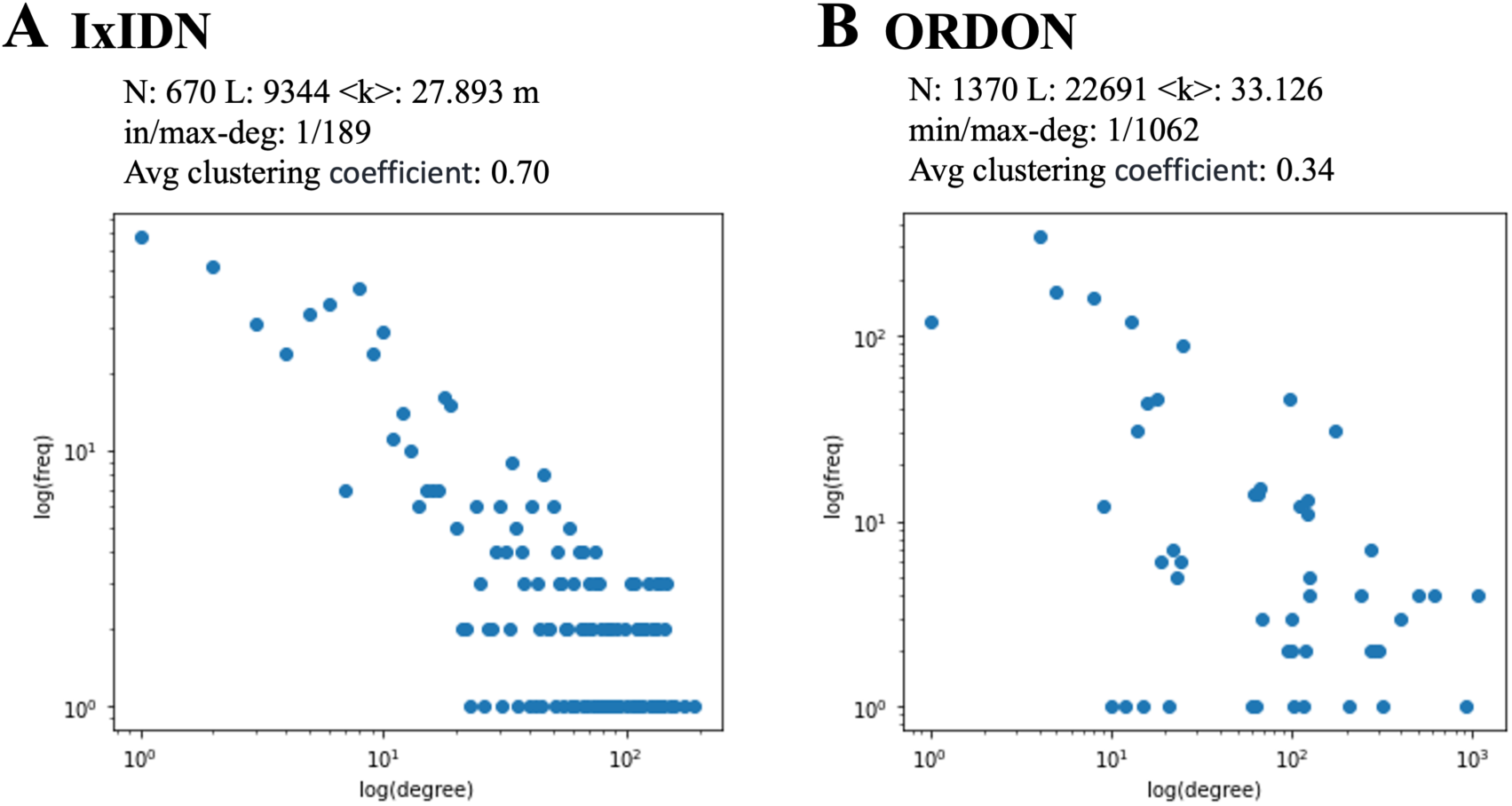
Degree Distributions of IxIDN and ORDON. Courtesy of Ravandi et al. in [40]. They exhibit majorly different patterns of interactions and structure compared to MechDist (Figure S5), making the transfer learning task from Dis2Vec to predict IxIDN and ORDON more difficult.

### S5. Compositionality of Dis2Vec Space

Compositionality is a characteristic of some well-structured vector representations [75] wherein analogical reasoning can be implemented as simple linear operations, like vector addition. In word representations for example, the closest word to the sum of “Russia” + “capital” might be “Moscow”. Some degree of compositionality is also evident in the Dis2Vec representations of diseases. Take the example of Mixed Connective Tissue Disease (ORPHA:809), which has overlapping features of the following 4 diseases in varying combinations: Systemic Lupus Erythematosus (ORPHA:536), Systemic Sclerosis (ORPHA:90291), Primary Sjogren Syndrome (ORPHA:289390), Systemic-onset juvenile idiopathic arthritis (ORPHA:85414), and Polymyositis (ORPHA:732), which are all in Cluster 31 although Polymyositis is located farther in the UMAP space.

Another example are the many paraneoplastic syndromes associated with Thymoma (ORPHA:99867), including Myasthenia gravis (ORPHA:589), and Dermatomyositis (ORPHA:221). The barycenter of these 2 diseases is most proximal Thymoma (after Myasthenia Gravis itself). These examples suggest that the positioning of overlap syndromes in the Dis2Vec representation corresponds roughly with linear combinations of disorders representing their overlapping features.

